# Comprehensive evolution and molecular characteristics of a large number of SARS-CoV-2 genomes revealed its epidemic trend and possible origins

**DOI:** 10.1101/2020.04.24.058933

**Authors:** Yunmeng Bai, Dawei Jiang, Jerome R Lon, Xiaoshi Chen, Meiling Hu, Shudai Lin, Zixi Chen, Xiaoning Wang, Yuhuan Meng, Hongli Du

## Abstract

**Objectives:** To reveal epidemic trend and possible origins of SARS-CoV-2 by exploring its evolution and molecular characteristics based on a large number of genomes since it has infected millions of people and spread quickly all over the world.

**Methods:** Various evolution analysis methods were employed.

**Results:** The estimated Ka/Ks ratio of SARS-CoV-2 is 1.008 or 1.094 based on 622 or 3624 SARS-CoV-2 genomes, and the time to the most recent common ancestor (tMRCA) was inferred in late September 2019. Further 9 key specific sites of highly linkage and four major haplotypes H1, H2, H3 and H4 were found. The Ka/Ks, detected population size and development trends of each major haplotype showed H3 and H4 subgroups were going through a purify evolution and almost disappeared after detection, indicating H3 and H4 might have existed for a long time, while H1 and H2 subgroups were going through a near neutral or neutral evolution and globally increased with time. Notably the frequency of H1 was generally high in Europe and correlated to death rate (r>0.37).

**Conclusions:** In this study, the evolution and molecular characteristics of more than 16000 genomic sequences provided a new perspective for revealing epidemiology of SARS-CoV-2.

## Introduction

The global outbreak of SARS-CoV-2 is currently and increasingly recognized as a serious, public health concern worldwide. Coronaviruses exist widely around the world, and to date, a total of seven types of coronaviruses that can infect humans have been found, of which four coronaviruses including hCoV-229E, hCoV-NL63, hCoV-OC43 and hCoV-HKU1 cause a cold, while the other three including SARS-CoV, MERS-CoV and SARS-CoV-2 usually cause mild to severe respiratory diseases. The total number of SARS-CoV and MERS-CoV infections are only 8,069 and 2,494 with reproduction number (R0) fluctuates 2.5-3.9 and 0.3-0.8, respectively. However, SARS-CoV-2 has infected 2471136 people in 212 countries up to April 22th(WHO, 2020), with the basic R0 ranging from 1.4 to 6.49(Liu, et al., 2020). Among these three typical coronavirus, MERS-CoV has the highest death rate of 34.40%, SARS-CoV has the modest death rate of 9.59%, SARS-CoV-2 is about 6.99% and 7.69% death rate of the global and Wuhan, but is quite high death rate in some countries of Europe, such as Belgium, Italy, United Kingdom, Netherlands, Spain and France, which even reaches 14.95%, 13.39%, 13.48%, 11.61%, 10.42% and 13.60% respectively according to the data of April 22th, 2020. Except for the shortage of medical supplies, aging and other factors, it is not clear whether there is virus mutation effect in these countries with such a significantly increased death rate.

It has been reported that SARS-CoV-2 belongs to beta-coronavirus and is mainly transmitted by the respiratory tract, which belongs to the same subgenus (SarbeCoVirus) as SARS-CoV(Lu, et al., 2020). Through analyzing the genome and structure of SARS-CoV-2, its receptor-binding domain (RBD) was found to bind with angiotensin-converting enzyme 2 (ACE2), which is also one of the receptors for binding SARS-CoV(Wrapp, et al., 2020). Some early genomic studies have shown that SARS-CoV-2 is similar to certain bat viruses (RaTG13, with the whole genome homology of 96.2%(Zhou, et al., 2020)) and Malayan pangolins coronaviruses (GD/P1L and GDP2S, with the whole genome homology of 92.4%(Lam, et al., 2020)). They have speculated several possible origins of SARS-CoV-2 based on its spike protein characteristics (cleavage sites or the RBD)(Lam, et al., 2020, Zhang and Holmes, 2020, Zhou, et al., 2020), in particular, SARS-CoV-2 exhibits 97.4% amino acid similarity to the Guangdong pangolin coronaviruses in RBD, even though it is most closely related to bat coronavirus RaTG13 at the whole genome level. However, it is not enough to present genome-wide evolution by a single gene or local evolution of RBD, whether bats or pangolins play an important role in the zoonotic origin of SARS-CoV-2 remains uncertain(Andersen, et al., 2020). Furthermore, since there is little molecular characteristics analysis of SARS-CoV-2 based on a large number of genomes, it is difficult to determine whether the virus has significant variation that affects its phenotype. Thereby, it is necessary to further reveal the phylogenetic evolution and molecular characteristics of the whole genome of SARS-CoV-2 in order to develop a comprehensive understanding of the virus and provide a basis for the prevention and treatment of SARS-CoV-2.

## Results

### Genome sequences

We got a total of 1053 genomic sequences up to March 22th, 2020. According to the filter criteria, 37 sequences with ambiguous time, 314 with low quality and 78 with similarity of 100% were removed. A total of 624 sequences were obtained to perform multiple sequences alignment. Two highly divergent sequences (EPI_ ISL_414690, EPI_ISL_415710) according to the firstly constructed phylogenetic tree were also filtered out (Table S1). The remaining 622 sequences were used to reconstruct a phylogenetic tree. In addition, total of 3624 and 16373 genomic sequences were redownloaded up to April 6th, 2020 and May 10th, 2020 respectively for further exploring the evolution and molecular characteristics of SARS-CoV-2.

### Estimate of evolution rate and the time to the most recent common ancestor for SARS-CoV, MERS-CoV, and SARS-CoV-2

In this study, date for SARS-CoV-2 ranged from 2019/12/26 to 2020/03/18 was collected. The average Ka/Ks of all the coding sequences was closer to 1 (1.008), which indicating the genome was going through a neutral evolution. We also reevaluated the Ka/Ks of SARS-CoV and MERS-CoV through the whole period, and found the ratio were smaller than SARS-CoV-2 (Table 1). To estimate more credible Ka/Ks for SARS-CoV-2, we recalculated it using redownloaded 3624 genome sequences ranged from 2019/12/26 to 2020/04/06, and the average Ka/Ks of it was 1.094 (Table 1), which was almost same with the above result.

**Table 1.**
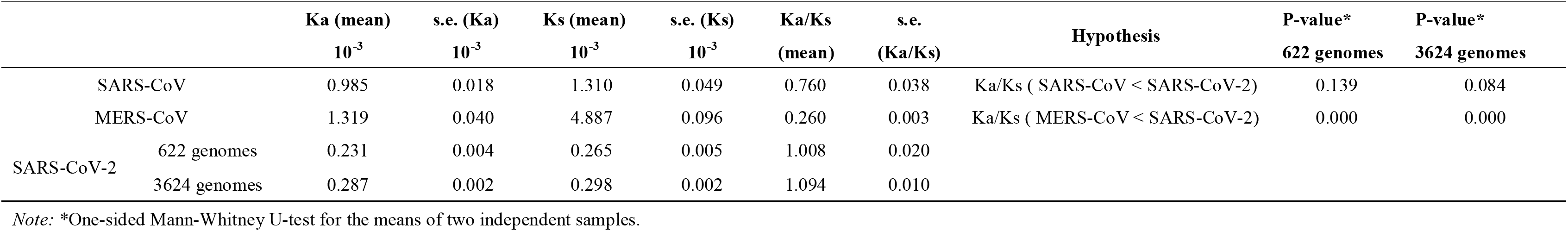
Statistics of Ka, Ks and Ka/Ks ratios for all coding regions of the SARS-CoV, MERS-CoV and SARS-CoV-2 genome sequences.

We assessed the temporal signal using TempEst v1.5.3(Rambaut, et al., 2016). All three datasets exhibit a positive correlation between root-to-tip divergence and sample collecting time (Figure S1), hence they are suitable for molecular clock analysis in BEAST(Bouckaert, et al., 2019, Rambaut, et al., 2016). The substitution rate of SARS-CoV-2 genome was estimated to be 1.601 × 10^−3^ (95% CI: 1.418 × 10^−3^ - 1.796 × 10^−3^, Table 2, Figure S2A) substitution/site/year, which is as in the same order of magnitude as SARS-CoV and MERS-CoV. The tMRCA was inferred on the late September, 2019 (95% CI: 2019/08/28-2019/10/26,Table 2, Figure S2B), about 2 months before the early cases of SARS-CoV-2(Huang, et al., 2020).

**Table 2.** Substitution rate and tMRCA estimated by BEAST v2.6.2.

|  | **substitution rate ($10^{-3}$)** | **95% CI ($10^{-3}$)** | **tMRCA** | **95% CI** | **references** |
| --- | --- | --- | --- | --- | --- |
| SARS-CoV | 1.050 | 0.489, 1.654 | 2002/6/28 | 2002/1/19, 2002/11/3 | this study |
|  | - | 0.800, 2.380 | spring of 2002 | - | <sup>17</sup> |
| MERS-CoV | 1.516 | 1.392, 1.632 | 2012/6/7 | 2012/6/4, 2012/6/9 | this study |
|  | 1.120 | 0.876, 1.370 | 2012/3 | 2011/12, 2012/6 | <sup>18</sup> |
| SARS-CoV-2 | 1.601 | 1.418, 1.796 | 2019/9/27 | 2019/8/28, 2019/10/26 | this study |

17. Zhao Z, Li H, Wu X, Zhong Y, Zhang K, Zhang YP, et al. Moderate mutation rate in the SARS coronavirus genome and its implications. BMC Evol Biol. 2004;4:21.
18. Cotten M, Watson SJ, Zumla AI, Makhdoom HQ, Palser AL, Ong SH, et al. Spread, circulation, and evolution of the Middle East respiratory syndrome coronavirus. mBio. 2014;5(1).

### Phylogenetic tree and clusters of SARS-CoV-2

The no-root phylogenetic trees constructed by the maximum likelihood method with PhyML 3.1(Guindon, et al., 2010) and MEGA(Kumar, et al., 2018) were showed in Figure 1, Figure S3A and 3B. According to the shape of phylogenetic trees, we divided 622 sequences into three clusters: Cluster 1 including 76 sequences mainly from North America, Cluster 2 including 367 sequences from all regions of the world, and Cluster 3 including 179 sequences mainly from Europe (Table S2).

**Figure 1.**
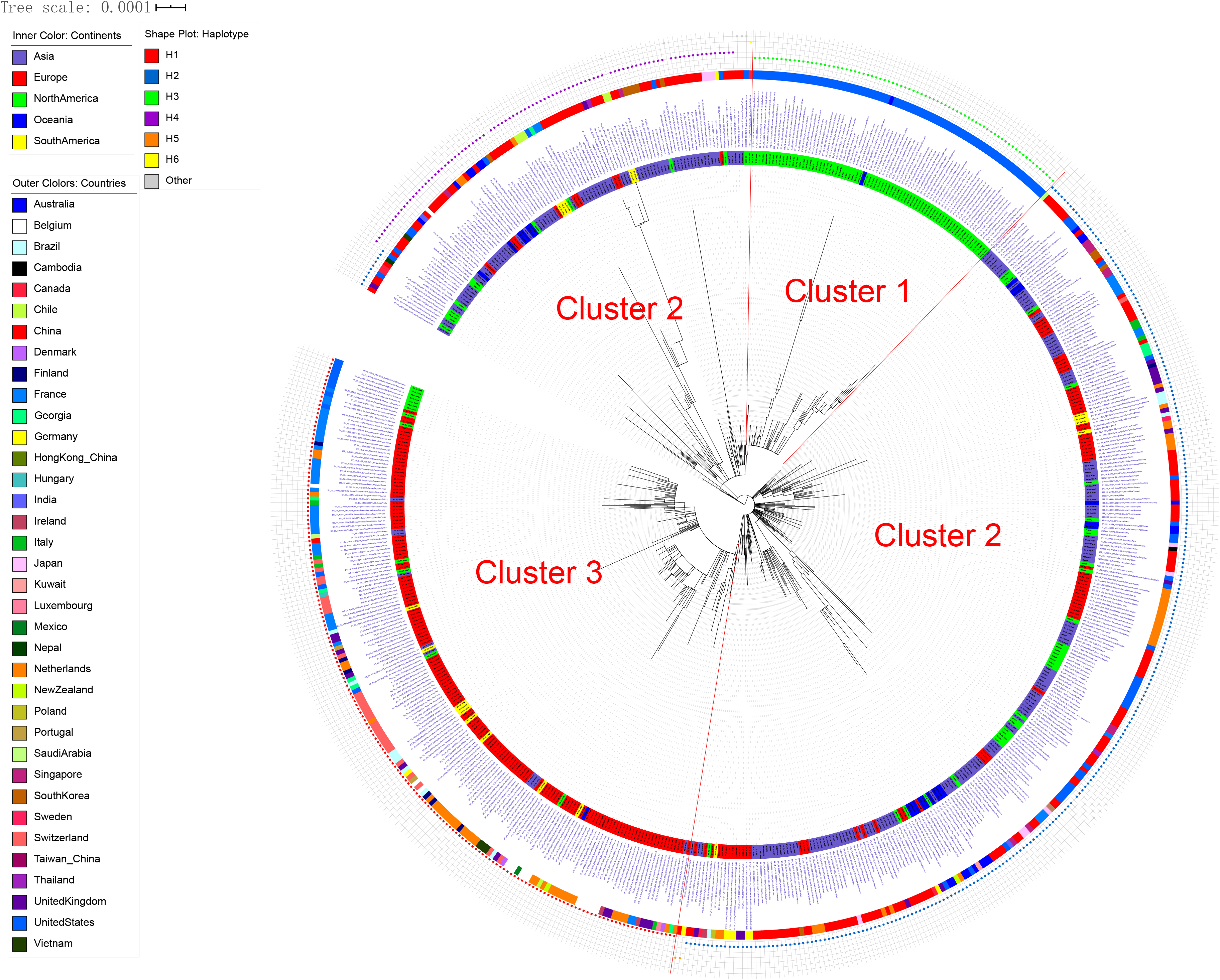
Phylogenetic tree and clusters of 622 SARS-CoV-2 genomes. The 622 sequences were clustered into three clusters: Cluster 1 were mainly from North America, Cluster 2 were from regions all over the world, and Cluster 3 were mainly from Europe.

### The specific sites of each Cluster

The *Fst* and population frequency of a total of 9 sites (NC_045512.2 as reference genome) were detected (Table 3, Table S3). Thereinto, three (C17747T, A17858G and C18060T) are the specific sites of Cluster 1, and four (C241T, C3037T, C14408T and A23403G) are the specific sites of Cluster 3. Notably, C241T was located in the 5’-UTR region and the others were located in coding regions (6 in *ofr1ab* gene, 1 in *S* gene and 1 in *ORF8* gene). Five of them were missense variant, including C14408T, C17747T and A17858G in *ofr1ab* gene, A23403G in *S* gene, and T28144C in *ORF8* gene. The PCA results showed that these 9 specific sites could clearly separate the three Clusters, while all SNV dataset could not clearly separate Cluster 1 and Cluster 2 (Figure S4), which further suggested that these 9 specific sites are the key sites for separating the three Clusters.

**Table 3.** The information of the 9 specific sites in each Cluster.

| Pos | Ref | Alt | Cluster 1 |  |  | Cluster 2 |  |  | Cluster 3 |  |  | Fst | Gene region | Mutation type | Protein changed | codon changed | Impact |
| --- | --- | --- | --- | --- | --- | --- | --- | --- | --- | --- | --- | --- | --- | --- | --- | --- | --- |
|  |  |  | pi | Major allele | frequency | pi | Major allele | frequency | pi | Major allele | frequency |  |  |  |  |  |  |
| 241 | C | T | 0.0000 | C | 1.0000 | 0.0109 | C | 0.9945 | 0.0000 | T | 1.0000 | 0.9912 | 5'UTR | upstream | NA | NA | MODIFIER |
| 3037 | C | T | 0.0000 | C | 1.0000 | 0.0109 | C | 0.9945 | 0.0000 | T | 1.0000 | 0.9912 | gene-orf1ab | synonymous | 924F | 2772ttC>ttT | LOW |
| 8782 | C | T | 0.0000 | T | 1.0000 | 0.3863 | C | 0.7390 | 0.0000 | C | 1.0000 | 0.5821 | gene-orf1ab | synonymous | 2839S | 8517agC>agT | LOW |
| 14408 | C | T | 0.0000 | C | 1.0000 | 0.0055 | C | 0.9973 | 0.0000 | T | 1.0000 | 0.9956 | gene-orf1ab | missense | 4715P>L | 14144cCt>cTt | MODERAT |
| 17747 | C | T | 0.0735 | T | 0.9620 | 0.0000 | C | 1.0000 | 0.0000 | C | 1.0000 | 0.9752 | gene-orf1ab | missense | 5828P>L | 17483cCt>cTt | MODERAT |
| 17858 | A | G | 0.0497 | G | 0.9747 | 0.0000 | A | 1.0000 | 0.0000 | A | 1.0000 | 0.9836 | gene-orf1ab | missense | 5865Y>C | 17594tAt>tGt | MODERAT |
| 18060 | C | T | 0.0000 | T | 1.0000 | 0.0164 | C | 0.9918 | 0.0000 | C | 1.0000 | 0.9761 | gene-orf1ab | synonymous | 5932L | 17796ctC>ctT | LOW |
| 23403 | A | G | 0.0000 | A | 1.0000 | 0.0164 | A | 0.9918 | 0.0000 | G | 1.0000 | 0.9869 | gene-S | missense | 614D>G | 1841gAt>gGt | MODERAT |
| 28144 | T | C | 0.0000 | C | 1.0000 | 0.3915 | T | 0.7335 | 0.0000 | T | 1.0000 | 0.5785 | gene-ORF8 | missense | 84L>S | 251tTa>tCa | MODERAT |

### Linkage of specific sites

We found the 9 specific sites are highly linkage based on 622 genome sequences (Figure 2A), then we carried out a further linkage analysis using the 3624 genome sequences (Figure 2B). As a result, for the 3624 genome sequences, 3 specific sites in the Cluster 1 were almost complete linkage, and haplotype CAC and TGT accounted for 98.65% of all the 3 site haplotypes. The same phenomenon was also found in 4 specific sites of Cluster 3, and haplotype CCCA and TTTG accounted for 97.68% of all the 4 site haplotypes. Intriguingly, the 9 specific sites were still highly linkage, and four haplotypes, including TTCTCACGT (H1), CCCCCACAT (H2), CCTCTGTAC (H3) and CCTCCACAC (H4), accounted for 95.89% of all the 9 site haplotypes. Thereinto, H1 and H3 had completely different bases at the 9 specific sites. The frequencies of each site and major haplotype for each country were showed in Figure 3 and Table S4. The data showed that the haplotype TTTG of the 4 specific sites in Cluster 3 had existed globally at present, and still exhibited high frequencies in most European countries but quite low in Asian countries. While the haplotype TGT of the 3 specific sites in Cluster 1 existed almost only in North America and Australia. For the 9 specific sites, most countries had only two or three major haplotypes, except America and Australia had all of four major haplotypes and with relative higher frequencies.

**Figure 2.**
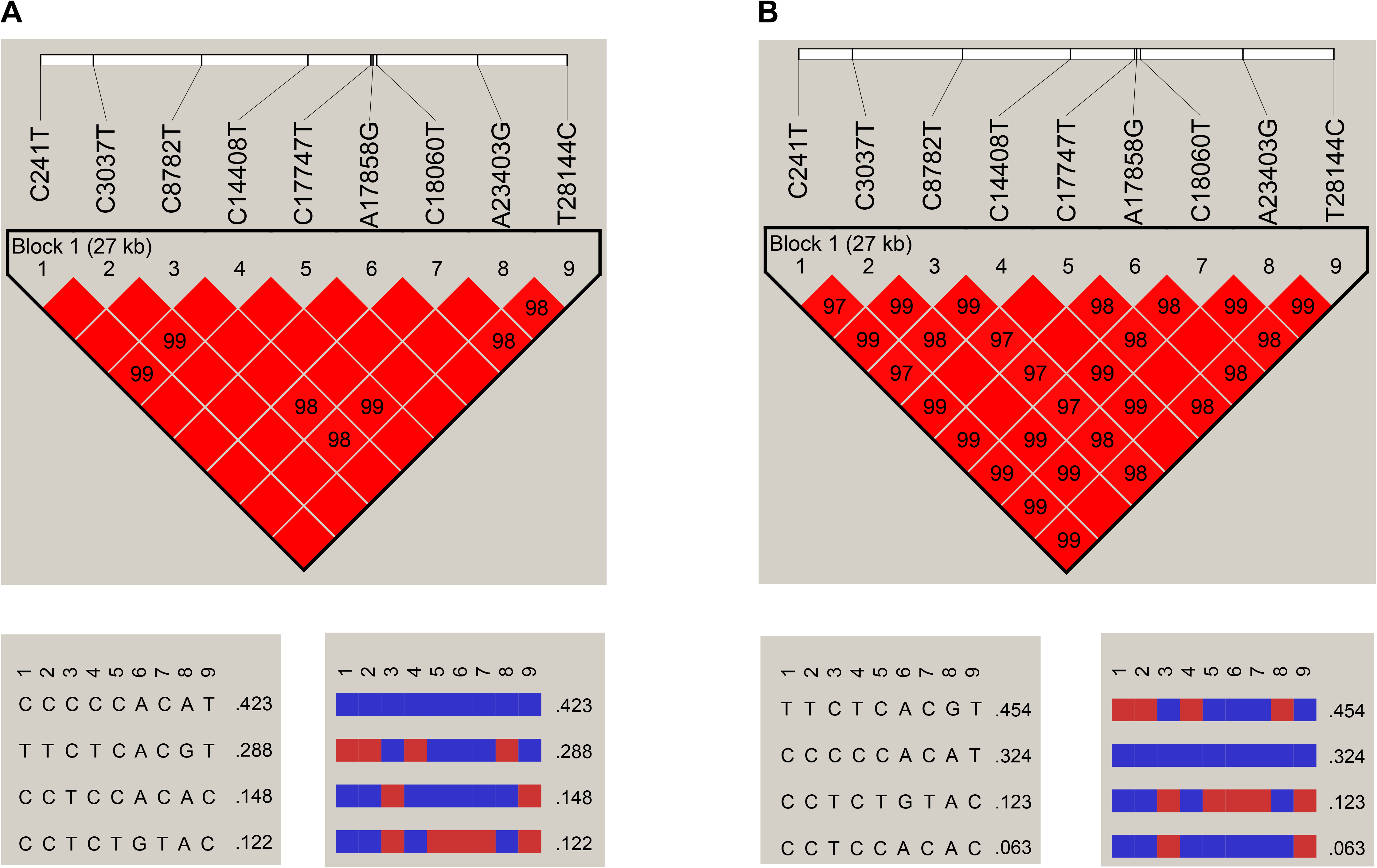
Linkage disequilibrium plot of haplotypes of the 9 specific sites. **A**. The plot for 622 genome sequences; **B**. The plot for 3624 genome sequences.

**Figure 3.**
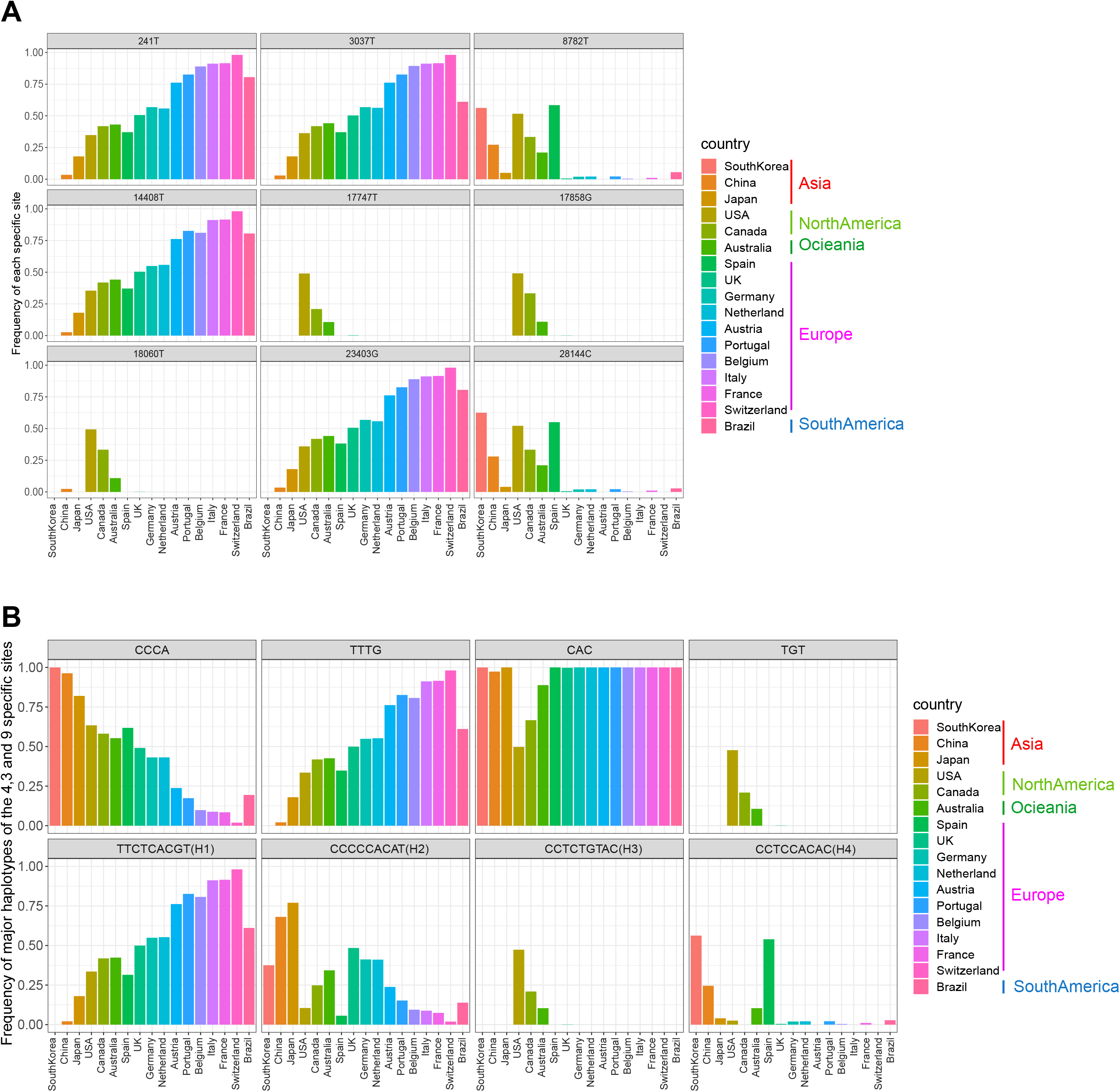
The frequencies of both the 9 specific sites and haplotypes. The frequencies of the 9 specific sites (**A**) and haplotypes (**B**) in each country for 3624 genomes.

### Characteristics and epidemic trends of major haplotype subgroups

All haplotypes of 9 specific sites for 3624 or 16373 genomes and the numbers of them were shown in Table S5. Four major haplotypes H1, H2, H3 and H4, three minor haplotypes H5, H7 and H8 close to H1 and one minor haplotype H6 close to H3 were found in both datasets. The numbers of these haplotypes for 16373 genomes with clear collection data detected in each country in chronological order were shown in Figure 4A. From these results, H2 and H4 haplotype subgroups had existed for a long time (2019-12-24 to 2020-04-28), the detected population size of H2 subgroup was far greater than that of H4 subgroup, and the H3 haplotype subgroup almost disappeared after detection (2020-02-18 to 2020-04-28), while H1 haplotype subgroup was globally increasing with time (2020-02-18 to 2020-05-05), which indicated H1 subgroup had adapted to the human hosts, and was under an adaptive growth period worldwide. However, due to the nonrandom sampling on early phase (only patients having a recent travel to Wuhan were detected), some earlier cases of H3 may be lost, the high proportion of H3 subgroup during February 18 to March 10 could confirm this point.

**Figure 4.**
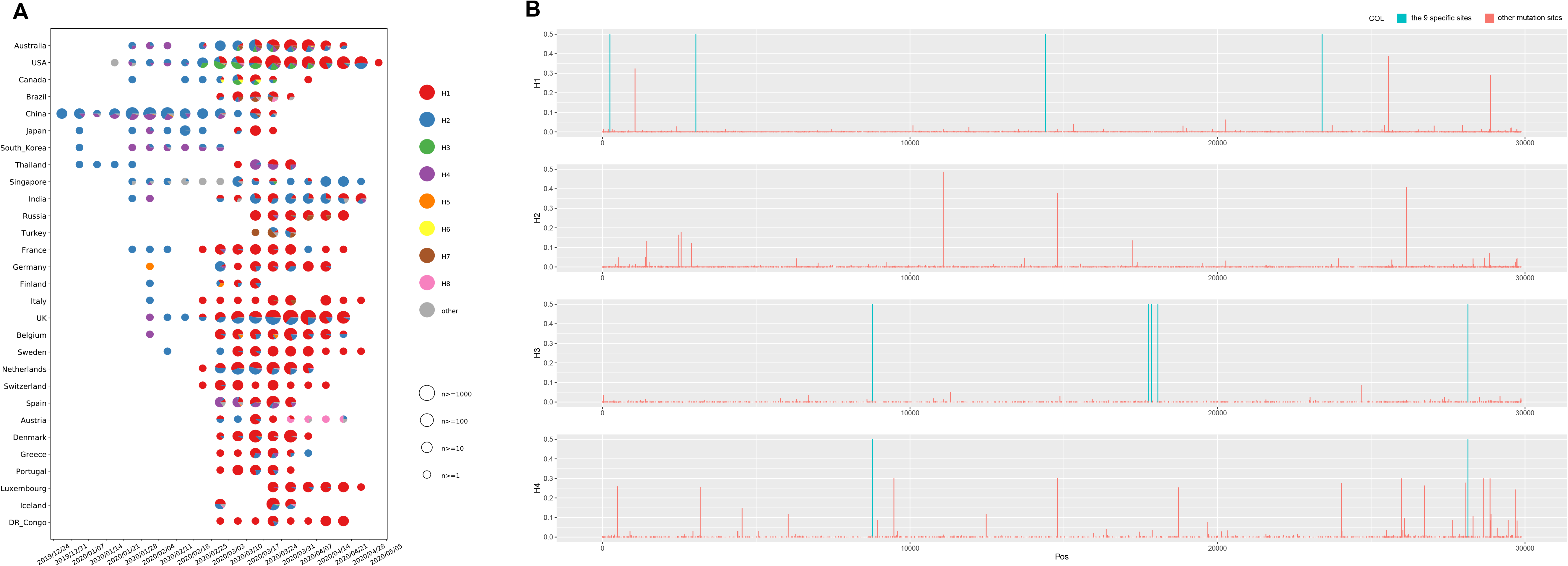
The characteristics of haplotype subgroups. **A**. The numbers of haplotypes of the 9 specific sites for 16373 genomes with clear collection data detected in each country in chronological order; **B**. The whole genome mutations in each major haplotype subgroup.

H3 and H4 subgroups had the lowest Ka/Ks ratio in 3624 or 16373 genomes among the 4 major subgroups (Table 4), suggesting that H3 and H4 subgroups might be going through a purifying evolution and have existed for a long time, while H2 and H1 subgroups might be going through a near neutral or neutral evolution according to their Ka/Ks ratios (Table 4), which were consistent with the above phenomenon that only H1 and H2 subgroups were expanding with time around the world. From the whole genome mutations in each major haplotype subgroup (Figure 4B, Table S6), we found that except the 9 specific sites, there was no common mutations with frequencies more than 0.05 in these haplotype subgroups but between H2 and H4 subgroups.

**Table 4.** Statistics of Ka, Ks and Ka/Ks ratios for all coding regions of each 4 major haplotype subgroup with 3624 and 16373 genomes respectively.

|  |  | **Ka (mean) 10<sup>-3</sup>** | **s.e. (Ka)10<sup>-3</sup>** | **Ks (mean) 10<sup>-3</sup>** | **s.e. (Ks)10<sup>-3</sup>** | **Ka/Ks (mean)** | **s.e. (Ka/Ks)** | **Hypothesis** | **P-value\*** |
| --- | --- | --- | --- | --- | --- | --- | --- | --- | --- |
| 3624 genomes | H1 | 0.510 | 0.057 | 0.498 | 0.056 | 1.099 | 0.010 | Ka/Ks ( H3 < H1) | 0.000 |
|  | H2 | 0.347 | 0.042 | 0.334 | 0.044 | 1.268 | 0.023 | Ka/Ks ( H3 < H2) | 0.000 |
|  | H3 | 0.298 | 0.007 | 0.397 | 0.009 | 0.795 | 0.010 | - | - |
|  | H4 | 0.334 | 0.023 | 0.414 | 0.019 | 0.796 | 0.019 | - | - |
| 16373 genomes | H1 | 0.416 | 0.011 | 0.384 | 0.011 | 1.178 | 0.005 | Ka/Ks ( H3 < H1) | 0.000 |
|  | H2 | 0.332 | 0.014 | 0.316 | 0.014 | 1.224 | 0.012 | Ka/Ks ( H3 < H2) | 0.000 |
|  | H3 | 0.305 | 0.004 | 0.401 | 0.005 | 0.809 | 0.007 | - | - |
|  | H4 | 0.354 | 0.010 | 0.477 | 0.009 | 0.749 | 0.010 | - | - |

*Note:* \*One-sided Mann-Whitney U-test for the means of two independent samples.

### Phylogenetic network of haplotype subgroups

Phylogenetic networks were inferred with 697 mutations called from 3624 genomes dataset. For these datasets, the network structures of TCS and MSN were similar. The major haplotype subgroups H4 and H2 were in the middle of the network, while H1 and H3 were in the end nodes of the network (Figure 5). According to the phylogenetic networks, we proposed four hypothesis: (1) the ancestral haplotypes evolved in four directions to obtain H1, H2, H3 and H4 respectively or evolved in two or more directions to obtain two or more major haplotypes and then involved into the other major haplotype(s); (2) H2 or H4 evolved in two directions respectively and finally generated H1 and H3; (3) H1 evolved in one direction to generate H2 and H4, and then evolved into H3; (4) H3 evolved in one direction to generate H4 and H2, and then evolved into H1. Although we cannot exclude the first hypothesis based on the present data, but if there are evolutionary relationships among the four major haplotypes, according to the Ka/Ks, detected population size and development trends in chronological order of each major haplotype subgroup, we speculate that the most likely evolution hypothesis is that H3 and H4 are the earliest haplotypes, which is gradually eliminated with selection, while H2 is the transitional haplotype in the evolution process, and H1 may be the haplotype to be finally fixed.

**Figure 5.**
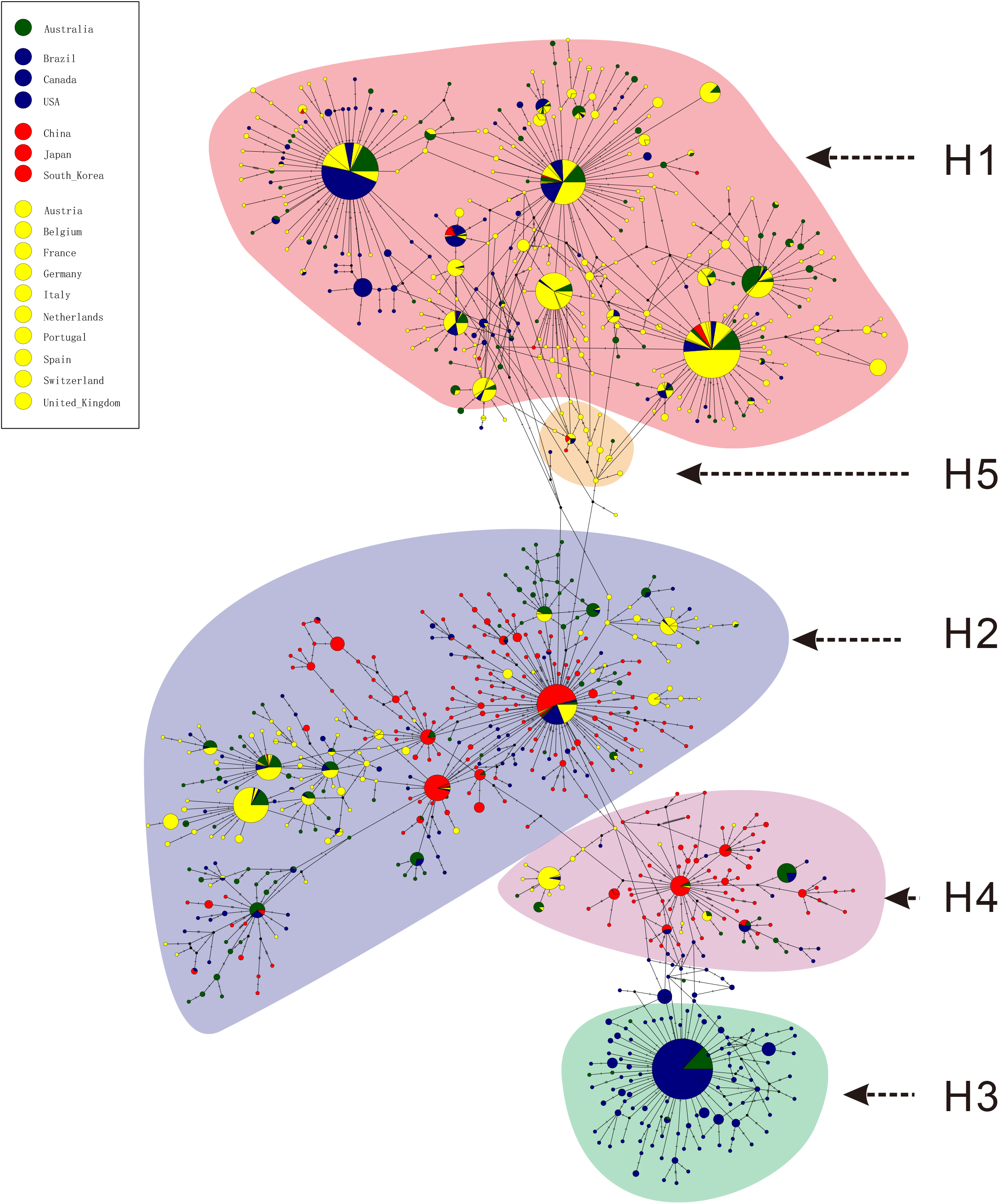
Phylogenetic network of haplotype subgroups for 3624 genomes. The network was inferred by POPART using TCS method. Each colored vertex represents a haplotype, with different colors indicating the different sampling areas. Hatch marks along edge indicated the number of mutations. Small black circles within the network indicated unsampled haplotypes. H1-H5 subgroups were point out according haplotypes of the 9 specific sites, and other small subgroups were not specially pointed out.

### Correlation analysis of specific sites with death rate and infectivity

To explore the relationship between death rate and the 9 specific sites, we used Pearson method to calculate the correlation coefficient between death rate and frequency of each specific site or major haplotype in 17 countries with 3624 genomes at early stage. As a result, all r values of 241T, 3037T, 14408T, 23403G and haplotype TTTG and H1 were more than 0.4 (Figure 6, Table S4). We also evaluated the correlation coefficient with 16373 genomes in 30 countries, the r values of haplotype TTTG and H1 were still more than 0.37 (Table S4), which might be increased when the death rates of some countries were further stabilized with time. These finding integrating their high frequencies in most European countries indicated that the 4 sites and haplotype TTTG and H1 might be related to the pathogenicity of SARS-CoV-2.

**Figure 6.**
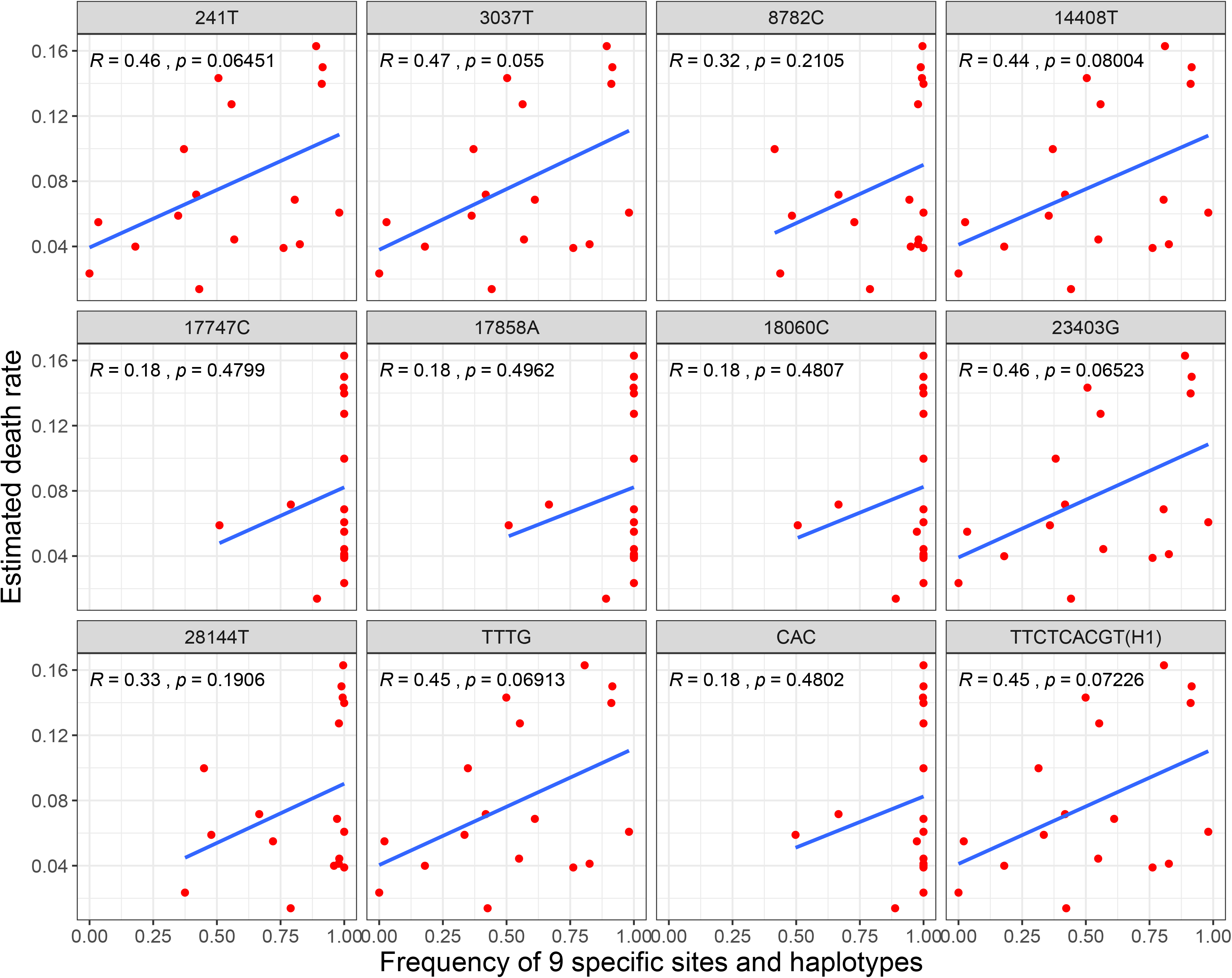
The correlation between death rate and frequencies of both the 9 specific sites and haplotypes.

To explore the relationship between infectivity and the 9 specific sites, we used population size of major haplotypes to deduce the possible specific sites related infectivity. We assumed that these major haplotype in each country were subject to similar virus transmission and control patterns, while the population sizes of H1 and H2 subgroups were far greater than those of H3 and H4 subgroups (Figure 4A, Table S4), then the common different specific sites of H1 and H2 subgroups with H3 and H4 subgroups, C8782T and T28144C, might be related to the infectivity of SARS-CoV-2, and the virus with C8782 and T28144 might be more infectious than those virus with 8782T and 28144C.

## Discussion

SARS-CoV-2 poses a great threat to the production, living and even survival of human beings(Guo, et al., 2020). With the further outbreak of SARS-CoV-2 in the world, further understanding on the evolution and molecular characteristics of SARS-CoV-2 based on a large number of genome sequences will help us cope better with the challenges brought by SARS-CoV-2.

Exploring evolution rate, tMRCA and phylogenetic tree of SARS-CoV-2 would help us better understand the virus(Yuen, et al., 2020). The average Ka/Ks for all the coding sequences of 622 and 3624 SARS-CoV-2 genomes was 1.008 and 1.094, which was higher than those of SARS-CoV and MERS-CoV, indicating that the SARS-CoV-2 was going through a neutral evolution. In general, SARS-CoV would show purifying selection pressures and the Ka/Ks would go down at the middle or end of the epidemic(He, et al., 2004). Therefore, it seemed to be reasonable considering that SARS-CoV-2 was under an exponential growth period around the world. Interestingly, we also found that the subgroups of different haplotypes (H1, H2, H3 and H4) seemed to experience different evolutionary patterns according to their Ka/Ks. The H3 haplotype subgroup disappeared soon after detection (2020-02-18 to 2020-04-28, Figure 4A), while H1 haplotype subgroup was globally increasing with time, these evolution and changes should be considered in developing therapeutic drugs and vaccines. The tMRCA of SARS-CoV-2 was inferred on the late September, 2019 (95% CI: 2019/08/28-2019/10/26), about 2 months before the early cases of SARS-CoV-2(Huang, et al., 2020). We also estimated tMRCA of SARS-CoV and MERS-CoV with the same methods, both were about 3 months later than the corresponding tMRCAs estimated by previous studied(Cotten, et al., 2014, Zhao, et al., 2004). A recent study used TreeDater method to estimate tMRCA for more than 7000 SARS-CoV-2 genomes and indicated the tMRCA of SARS-CoV-2 was around 6 October 2019 to 11 December 2019, which were about one and a half months later than ours and in broad agreement with six previous studies all performed on no more than 120 early SARS-CoV-2 genomes with BEAST method(van Dorp, et al., 2020). While we used the most common method BEAST to estimate tMRCA of SARS-CoV-2 based on 622 genomes and combined with the comparison of tMRCA of SARS-CoV and MERS-CoV, indicating that the estimated tMRCA in the present study are more reliable.

A recent study clustered 160 SARS-CoV-2 whole-genome sequences into A, B and C groups by a phylogenetic network analysis by taking bat RaTG13 as root (Forster, et al., 2020). The cluster result was similar to our study: both the samples in Cluster A and our Cluster 1 were mainly from the United States; the samples in Cluster C and our Cluster 3 were mainly from European countries, while Cluster B and our Cluster 2 were mainly from China and the other regions. It was interesting that the markers C8782T and T28144C, which were also discovered by Yu et al(Yu, et al., 2020), in Cluster B were also found in our study, but the other markers in Cluster A (T29095C) and Cluster C (G26144T) were not significantly in our study. That may be caused by different sample sizes and different constructing methods of phylogenetic tree. Based on the base substitution model, the ML method avoids the possible “long-branch attraction” problem in the maximum parsimony method and is faster than the Bayesian method(Holder and Lewis, 2003), hence it could be used as a reliable method for phylogenetic analysis. Some studies used the genome of bat SARS-like-CoV(Zhang, et al., 2020), RaTG13(Zhang, et al., 2020) or MT019529 (https://bigd.big.ac.cn/ncov/tree) as the root of phylogenetic tree. Unfortunately, there was no obvious evidence showing that SARS-CoV-2 was from the bat coronavirus even the identity between SARS-CoV-2 and RaTG13 was up to 96.2%(Zhou, et al., 2020). In our study, the tMRCA of SARS-CoV-2 was inferred on the late September 2019, which indicated there might exist an earlier SARS-CoV-2 strain we didn’t find. Then in the case of unclear source of SARS-CoV-2 and high homology of its genomes (> 99.9% homology mostly), it may be inappropriate to identify the evolutionary characteristics inside the genomes by taking bat SARS-like-CoV, RaTG13 or MT019529 as root. Therefore, the maximum likelihood method was used in current study to construct a no-root tree to obtain the reliable clusters with different characteristics. Based on the no-root tree, we identified 9 specific sites of highly linkage that play a decisive role in the classification of clusters successfully. Among the four major haplotypes, H1 and H3 were in Cluster 3 and Cluster 1, respectively, while there were two haplotypes H2 and H4 in Cluster 2 (Figure 1, Figure S3B).

Among the 9 specific sites, 8 of them were located in coding regions (6 in *ofr1ab* gene, 1 in *S* gene and 1 in *ORF8* gene), and 5 of them were missense variant, including C14408T, C17747T and A17858G in *ofr1ab* gene, A23403G in *S* gene, and C28144T in *ORF8* gene. *Orf1ab* is composed of two partially overlapping open reading frames (orf), orf1a and 1b. It is proteolytic cleaved into 16 non-structural proteins (nsp), including nsp1 (suppress antiviral host response), nsp9 (RNA/DNA binding activity), nsp12 (RNA-dependent RNA polymerase), nsp13 (helicase) and others(Chan, et al., 2020), indicating the vital role of it in transcription, replication, innate immune response and virulence(Graham, et al., 2008). C14408T with high frequencies of T in European countries were located at nsp12 region, indicating that this missense variant might influence the role of RNA polymerase. Spike glycoprotein, the largest structural protein on the surface of coronaviruses, comprises of S1 and S2 subunits which mediating binding the receptor on the host cell surface and fusing membranes, respectively(Li, 2016). It has been reported that S protein of SARS-CoV-2 can bind angiotensin-converting enzyme 2 (ACE2) with higher affinity than that of SARS(Wrapp, et al., 2020, Zhou, et al., 2020). Whether the missense variant of A23403G in *S* gene, which with high frequencies of G in European countries, will change the affinity between S protein and ACE2 remains to be further investigated.

It seems to take a long time to finally fix mutations according to the mutation frequency of each subgroup. For example, H2 and H4 subgroups, which have been detected for more than four months from December 24, 2019 to May 5, 2020 (Figure 4A), have more mutations with higher frequencies, but the highest mutation frequency is only 0.486 at the position of 11083 (Figure 4B, Table S6). From these phenomena, it can be inferred that it takes a long time for the specific sites of each major subgroup to be fixed, but it may be faster if the early population is small. In addition, there is also the possibility that an ancestor strain evolved in four directions by obtaining the specific mutations directly and produced the four current major haplotypes, so the evolution time for obtaining four major haplotypes may be shorter, which seems to be consistent with the phenomenon that four major haplotypes can be detected in two months (Figure 4A). However, if there exist evolution relationship among the major haplotypes of SARS-CoV-2, it is difficult to complete the evolution among the four major haplotypes within two months (2019-12-24 to 2020-02-18, Figure 4A) at the current evolution rate of each major haplotype population (Table 4). Therefore, we speculate that the transformation among the four major haplotypes may have been completed for a long time, which have not been detected. What’s interesting is that only United States and Australia among 29 countries had all of four major haplotypes and with relative higher frequency (Figure 4A, Table S4), which indicated that the two countries are the most likely places where the virus appeared earlier based on all the present data.

Previous studies showed that the death rate of SARS-CoV-2 is affected by many factors, such as medical supplies, aging, life style and other factors(Liu, et al., 2020, Malavolta, et al., 2020). However, our study found that the 4 specific sites (C241T, C3037T, C14408T and A23403G) in the Cluster 3 were almost complete linkage, and the frequency of haplotype TTTG was generally high in European countries and correlated to death rate (r>0.37) both based on 3624 or 16373 SARS-CoV-2 genomes, which provides a new perspective to the reasons of relatively high death rate in Europe, and these specific sites should be considered in designing new vaccine and drug development of SARS-CoV-2. Two possible specific sites C8782T and T28144C related to the infectivity of SARS-CoV-2 were also deduced in the present study, which would provide some basis for SARS-CoV-2 epidemiology.

## Conclusion and prospective

The Ka/Ks ratio of SARS-CoV-2 and tMRCA of SARS-CoV-2 (95% CI: 2019/08/28-2019/10/26) indicated that SARS-CoV-2 might have completed the selection pressure of cross-host evolution in the early stage and be going through a neutral evolution at present. The 9 specific sites with highly linkage were found to play a decisive role in the classification of clusters. Thereinto, 3 of them are the specific sites of Cluster 1 mainly from North America and Australia, 4 of them are the specific sites of Cluster 3 mainly from Europe in the early stage, both of these 3 and 4 specific sites are almost complete linkage. The frequencies of haplotype TTTG for the 4 specific sites and H1 for 9 specific sites were generally high in European countries and correlated to death rate (r>0.37) based on 3624 or 16373 SARS-CoV-2 genomes, which suggested these haplotypes might relate to pathogenicity of SARS-CoV-2 and provided a new perspective to the reasons of relatively high death rates in Europe. The relationship between the haplotype TTTG or H1 and the pathogenicity of SARS-CoV-2 needs to be further verified by clinical samples or virus virulence test experiment, and the 9 specific sites with highly linkage may provide a starting point for the traceability research of SARS-CoV-2. Given that the different evolution patterns of different haplotypes subgroups, we should consider these evolution and changes in the development of therapeutic drugs and vaccines.

## Materials and methods

### Genome sequences

The complete genome sequences of SARS-CoV-2 were downloaded from China National Center for Bioinformation (https://bigd.big.ac.cn/ncov/release_genome/) and GISAID (https://www.gisaid.org/) up to March 22th, 2020. The sequences were filtered out according to the following criteria: (1) sequences with ambiguous time; (2) sequences with low quality which contained the counts of unknown bases > 15 and degenerate bases > 50 (https://bigd.big.ac.cn/ncov/release_genome); (3) sequences with similarity of 100% were removed to unique one. Finally, 624 high quality genomes with precise collection time were selected and aligned using MAFFT v7 with automatic parameters. Besides, the genome sequences of 7 SARS-CoV and 475 MERS-CoV were also downloaded from NCBI (https://www.ncbi.nlm.nih.gov/), and the MERS-CoV dataset including samples collected from both human and camel. In addition, for further exploring evolution and molecular characteristics of SARS-CoV-2 based on the larger amount of genomic data, we redownloaded validation datasets of the genome sequences from GISAID up to April 6th, 2020 and May 10th, 2020 respectively.

### Estimate of evolution rate and the time to the most recent common ancestor for SARS-CoV, MERS-CoV, and SARS-CoV-2

The average Ka, Ks and Ka/Ks for all coding sequences were calculated using KaKs_Calculator v1.2(Zhang, et al., 2006), and the substitution rate and tMRCA were estimated using BEAST v2.6.2(Bouckaert, et al., 2019). The temporal signal with root-to-tip divergence was visualized in TempEst v1.5.3(Rambaut, et al., 2016) using a ML whole genome tree with bootstrap value as input. For SARS-CoV and SARS-CoV-2, we selected a strict molecular clock and Coalescent Exponential Population Model. For MERS-CoV, we selected a relaxed molecular clock and Birth Death Skyline Serial Cond Root Model. We used the tip dates and chose the HKY as the site substitution model in all these analyses. Markov Chain Monte Carlo (MCMC) chain length was set to 10,000,000 steps sampling after every 1000 steps. The output was examined in Tracer v1.6 (http://tree.bio.ed.ac.uk/software/tracer/).

### Variants calling of SARS-CoV-2 genome sequences

Each genome sequence was aligned to the reference genome (NC_045512.2) using bowtie2 with default parameters(Langmead and Salzberg, 2012), and variants were called by samtools (sort; mpileup -gf) and bcftoots (call -vm). The merge VCF files were created by bgzip and bcftools (merge --missing-to-ref)(Li, 2011, Li, et al., 2009).

### Phylogenetic tree construction and virus isolates clustering for SARS-CoV-2

After alignment of 624 SARS-CoV-2 genomes with high quality and manually deleted 2 highly divergent genomes (EPI_ISL_415710, EPI_ISL_414690) according to the firstly constructed phylogenetic tree, the aligned dataset of 622 sequences was phylogenetically analyzed. The SMS method was used to select GTR+G as the base substitution model(Lefort, et al., 2017), and the PhyML 3.1(Guindon, et al., 2010) and MEGA(Kumar, et al., 2018) were used to construct the no-root phylogenetic tree by the maximum likelihood method with the bootstrap value of 100. The online tool iTOL(Letunic and Bork, 2019) was used to visualized the phylogenetic tree. The clusters were defined by the shape of phylogenetic tree.

### Detection of specific sites from each Cluster

Information (ID, countries/regions and collection time) and variants (NC_045512.2 as reference genome) of each genome from each Cluster were extracted. The allele frequency and nucleotide divergency (pi) for each site in the virus population of each Cluster were measured by vcftools(Danecek, et al., 2011). The *Fst* were also calculated by vcftools(Danecek, et al., 2011) to assess the diversity between the Clusters. Sites with high level of *F*st together with different major allele in each Cluster were filtered as the specific sites. PCA were analyzed by the GCTA v1.93.1beta(Yang, et al., 2011) with the specific sites and all SNV dataset respectively.

### Linkage analysis of specific sites and characteristics of major haplotype subgroups

The linkage disequilibrium of the specific sites were analyzed by haploview(Barrett, et al., 2005), and the statistics of the haplotype of the specific sites for each Cluster or country were used in-house perl script.

### Phylogenetic network of haplotype subgroups

The phylogenetic networks were inferred by PopART package v1.7.2(Leigh, et al., 2015) using TCS and minimum spanning network (MSN) methods respectively.

### Frequencies of specific sites or haplotypes and correlation with death rate

The frequencies of specific sites for each country were calculated. The death rate was estimated with Total Deaths /Confirmed Cases based on the data from Johns Hopkins resources on May 12th, 2020 (https://www.arcgis.com/apps/opsdashboard/index.html#/bda7594740fd40299423467b48e9ecf6).The correlation coefficient between death rate and frequencies of specific site or haplotype in different countries was calculated using Pearson method.

## Authors’ contributions

HD conceived the study. YM, YB, DJ, JL and XC carried out the data analysis and wrote the manuscript. MH, SL, and ZC collected data and revised the manuscript. XW attended the discussions. HD and YM supervised the whole work and revised the manuscript.

## Ethical Approval

Not required.

## Supplementary material

**Figure S1 Regression analyses of the root-to-tip divergence with the maximum-likelihood phylogenetic tree of SARS-CoV (A), MERS-CoV (B) and SARS-CoV-2 (C)**

The root-to-tip genetic distances were inferred in TempEst v1.5.3. Three data sets exhibit a positive correlation between root-to-tip divergence and sample collecting time, and they appear to be suitable for molecular clock analysis.

**Figure S2 Substitution rate and tMRCA estimate of SARS-CoV-2**

**A**. Estimated substitution rate of SARS-CoV-2 using BEAST v2.6.2 under a strict molecular clock. The mean rate was 1.601 × 10^−3^ (substitutions per site per year). **B**. The estimated tMRCA of SARS-CoV-2 using BEAST v2.6.2 under a strict molecular clock. The mean tMRCA was 2019.74 (date: 2019-09-27).

**Figure S3 Phylogenetic tree of 622 SARS-CoV-2 genomes**

**A**. Phylogenetic tree constructed by PhyML with bootstrap value of 100; **B**. Phylogenetic tree constructed by MEGA with bootstrap value of 100.

**Figure S4 The PCAs analysis result based on the 9 specific sites (A) and all SNVs (B)**

**Table S1 The information of the first downloaded complete genome sequences**

**Table S2 The sequence IDs in each Cluster**

**Table S3 The details information of the SNVs in all Clusters**

**Table S4 The information of death rate, frequencies of the 9 specific sites and major haplotypes in each country for 3624 and 16373 genomes respectively**

**Table S5 All haplotypes of the 9 specific sites and numbers of them for 3624 and 16373 genomes respectively**

**Table S6 Whole genome mutations and frequencies in each major haplotype subgroup of 3624 and 16373 genomes respectively**

## References

Andersen KG, Rambaut A, Lipkin WI, Holmes EC, Garry RF. (2020) The proximal origin of SARS-CoV-2. Nature Medicine, 10.1038/s41591-020-0820-9

Barrett JC, Fry B, Maller J, Daly MJ. (2005) Haploview: analysis and visualization of LD and haplotype maps. Bioinformatics, 21:263–5.

Bouckaert R, Vaughan TG, Barido-Sottani J, et al. (2019) BEAST 2.5: An advanced software platform for Bayesian evolutionary analysis. PLoS Comput Biol, 15:e1006650.

Chan JF, Kok KH, Zhu Z, et al. (2020) Genomic characterization of the 2019 novel human-pathogenic coronavirus isolated from a patient with atypical pneumonia after visiting Wuhan. Emerg Microbes Infect, 9:221–36.

Cotten M, Watson SJ, Zumla AI, et al. (2014) Spread, circulation, and evolution of the Middle East respiratory syndrome coronavirus. mBio, 5:

Danecek P, Auton A, Abecasis G, et al. (2011) The variant call format and VCFtools. Bioinformatics (Oxford, England), 27:2156–8.

Forster P, Forster L, Renfrew C, Forster M. (2020) Phylogenetic network analysis of SARS-CoV-2 genomes. Proceedings of the National Academy of Sciences, 10.1073/pnas.2004999117202004999.

Graham RL, Sparks JS, Eckerle LD, Sims AC, Denison MR. (2008) SARS coronavirus replicase proteins in pathogenesis. Virus Res, 133:88–100.

Guindon S, Dufayard JF, Lefort V, Anisimova M, Hordijk W, Gascuel O. (2010) New algorithms and methods to estimate maximum-likelihood phylogenies: assessing the performance of PhyML 3.0. Syst Biol, 59:307–21.

Guo YR, Cao QD, Hong ZS, et al. (2020) The origin, transmission and clinical therapies on coronavirus disease 2019 (COVID-19) outbreak - an update on the status. Mil Med Res, 7:11.

He J, Peng G, Min J. (2004) Molecular Evolution of the SARS Coronavirus During the Course of the SARS Epidemic in China. Science, 303:

Holder M, Lewis PO. (2003) Phylogeny estimation: traditional and Bayesian approaches. Nat Rev Genet, 4:275–84.

Huang C, Wang Y, Li X, et al. (2020) Clinical features of patients infected with 2019 novel coronavirus in Wuhan, China. Lancet, 395:497–506.

Kumar S, Stecher G, Li M, Knyaz C, Tamura K. (2018) MEGA X: Molecular Evolutionary Genetics Analysis across Computing Platforms. Mol Biol Evol, 35:1547–9.

Lam TT, Shum MH, Zhu HC, et al. (2020) Identifying SARS-CoV-2 related coronaviruses in Malayan pangolins. Nature, 10.1038/s41586-020-2169-0

Langmead B, Salzberg SL. (2012) Fast gapped-read alignment with Bowtie 2. Nat Methods, 9:357–9.

Lefort V, Longueville JE, Gascuel O. (2017) SMS: Smart Model Selection in PhyML. Mol Biol Evol, 34:2422–4.

Leigh JW, Bryant D, Nakagawa S. (2015) popart: full‐feature software for haplotype network construction. Methods in Ecology and Evolution, 6:1110–6.

Letunic I, Bork P. (2019) Interactive Tree Of Life (iTOL) v4: recent updates and new developments. Nucleic Acids Res, 47:W256–W9.

Li F. (2016) Structure, Function, and Evolution of Coronavirus Spike Proteins. Annu Rev Virol, 3:237–61.

Li H. (2011) A statistical framework for SNP calling, mutation discovery, association mapping and population genetical parameter estimation from sequencing data. Bioinformatics (Oxford, England), 27:2987–93.

Li H, Handsaker B, Wysoker A, et al. (2009) The Sequence Alignment/Map format and SAMtools. Bioinformatics (Oxford, England), 25:2078–9.

Liu Y, Du X, Chen J, et al. (2020) Neutrophil-to-lymphocyte ratio as an independent risk factor for mortality in hospitalized patients with COVID-19. J Infect, 10.1016/j.jinf.2020.04.002

Liu Y, Gayle AA, Wilder-Smith A, Rocklov J. (2020) The reproductive number of COVID-19 is higher compared to SARS coronavirus. J Travel Med, 27:

Lu R, Zhao X, Li J, et al. (2020) Genomic characterisation and epidemiology of 2019 novel coronavirus: implications for virus origins and receptor binding. Lancet, 395:565–74.

Malavolta M, Giacconi R, Brunetti D, Provinciali M, Maggi F. (2020) Exploring the Relevance of Senotherapeutics for the Current SARS-CoV-2 Emergency and Similar Future Global Health Threats. Cells, 9:

Rambaut A, Lam TT, Max Carvalho L, Pybus OG. (2016) Exploring the temporal structure of heterochronous sequences using TempEst (formerly Path-O-Gen). Virus Evol, 2:vew007.

van Dorp L, Acman M, Richard D, et al. (2020) Emergence of genomic diversity and recurrent mutations in SARS-CoV-2. Infection, Genetics and Evolution, https://doi.org/10.1016/j.meegid.2020.104351104351.

WHO. (2020) Coronavirus disease 2019 (COVID-19) Situation Report – 93. WHO,

Wrapp D, Wang N, Corbett KS, et al. (2020) Cryo-EM structure of the 2019-nCoV spike in the prefusion conformation. Science, 367:1260–3.

Yang J, Lee SH, Goddard ME, Visscher PM. (2011) GCTA: a tool for genome-wide complex trait analysis. Am J Hum Genet, 88:76–82.

Yu WB, Tang GD, Zhang L, Corlett RT. (2020) Decoding the evolution and transmissions of the novel pneumonia coronavirus (SARS-CoV-2 / HCoV-19) using whole genomic data. Zool Res, 41:247–57.

Yuen K-S, Ye Z-W, Fung S-Y, Chan C-P, Jin D-Y. (2020) SARS-CoV-2 and COVID-19: The most important research questions. Cell Biosci, 10:40.

Zhang L, Shen F-m, Chen F, Lin Z. (2020) Origin and Evolution of the 2019 Novel Coronavirus. Clinical Infectious Diseases, 10.1093/cid/ciaa112

Zhang YZ, Holmes EC. (2020) A Genomic Perspective on the Origin and Emergence of SARS-CoV-2. Cell, 181:223–7.

Zhang Z, Li J, Zhao XQ, Wang J, Wong GK, Yu J. (2006) KaKs_Calculator: calculating Ka and Ks through model selection and model averaging. Genomics Proteomics Bioinformatics, 4:259–63.

Zhao Z, Li H, Wu X, et al. (2004) Moderate mutation rate in the SARS coronavirus genome and its implications. BMC Evol Biol, 4:21.

Zhou P, Yang XL, Wang XG, et al. (2020) A pneumonia outbreak associated with a new coronavirus of probable bat origin. Nature, 579:270–3.

