## Supplementary figures and images for "Comprehensive evolution and molecular characteristics of a large number of SARS-CoV-2 genomes revealed its epidemic trend and possible origins"

### Fig S1

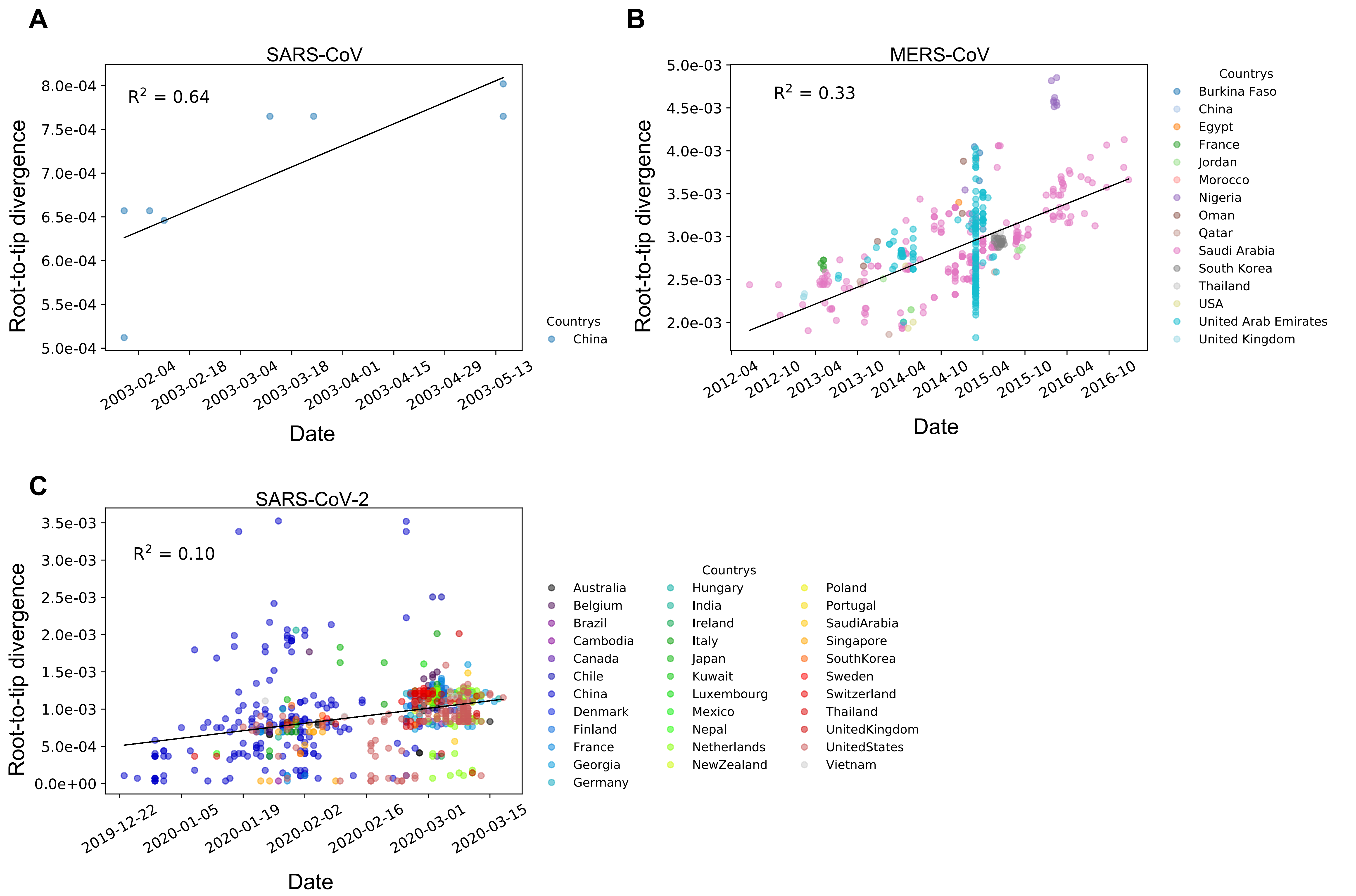

### Fig S2

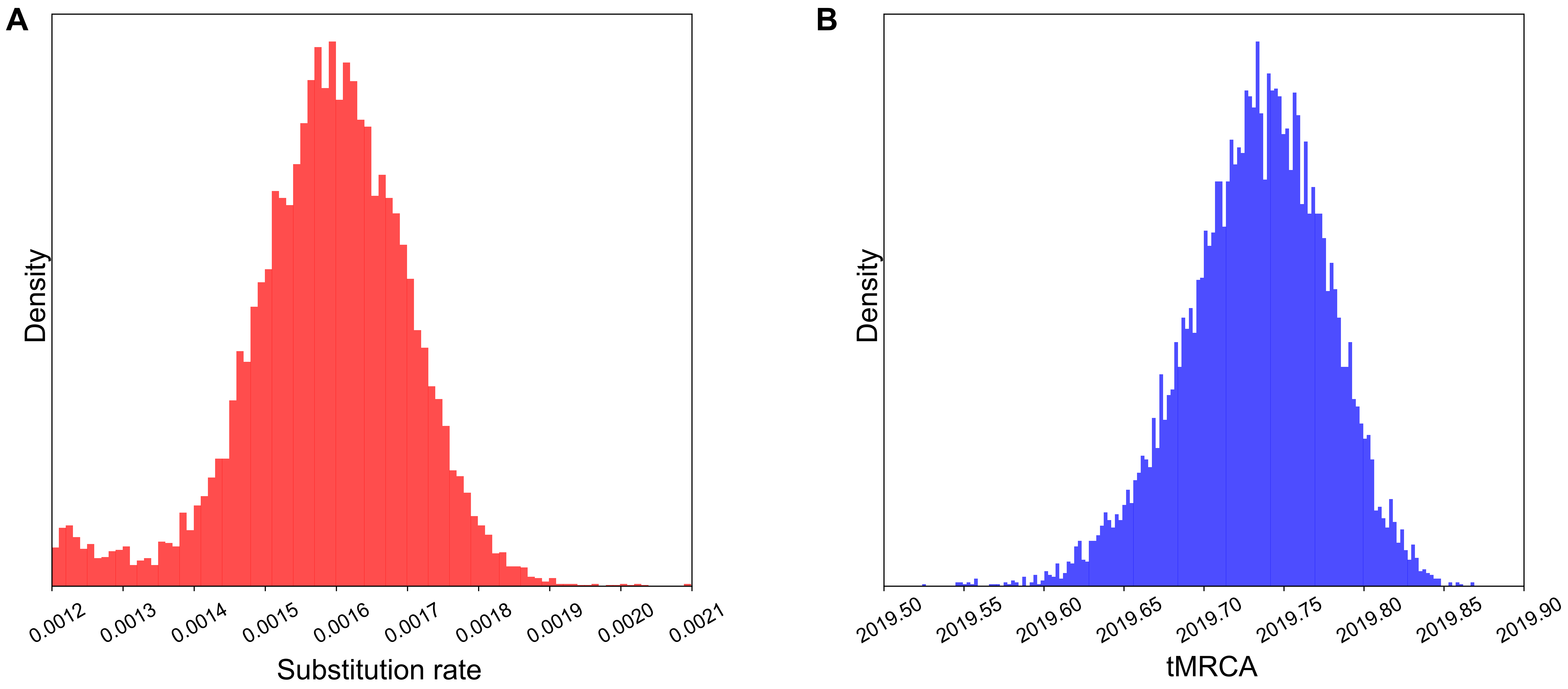

### Fig S3

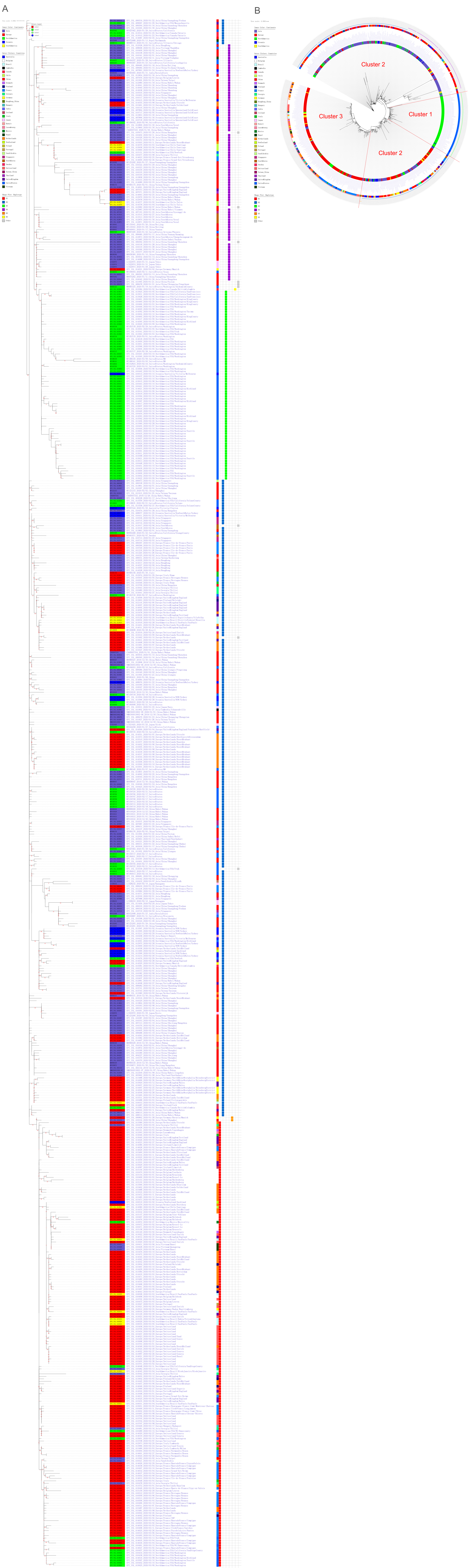

### Fig S4

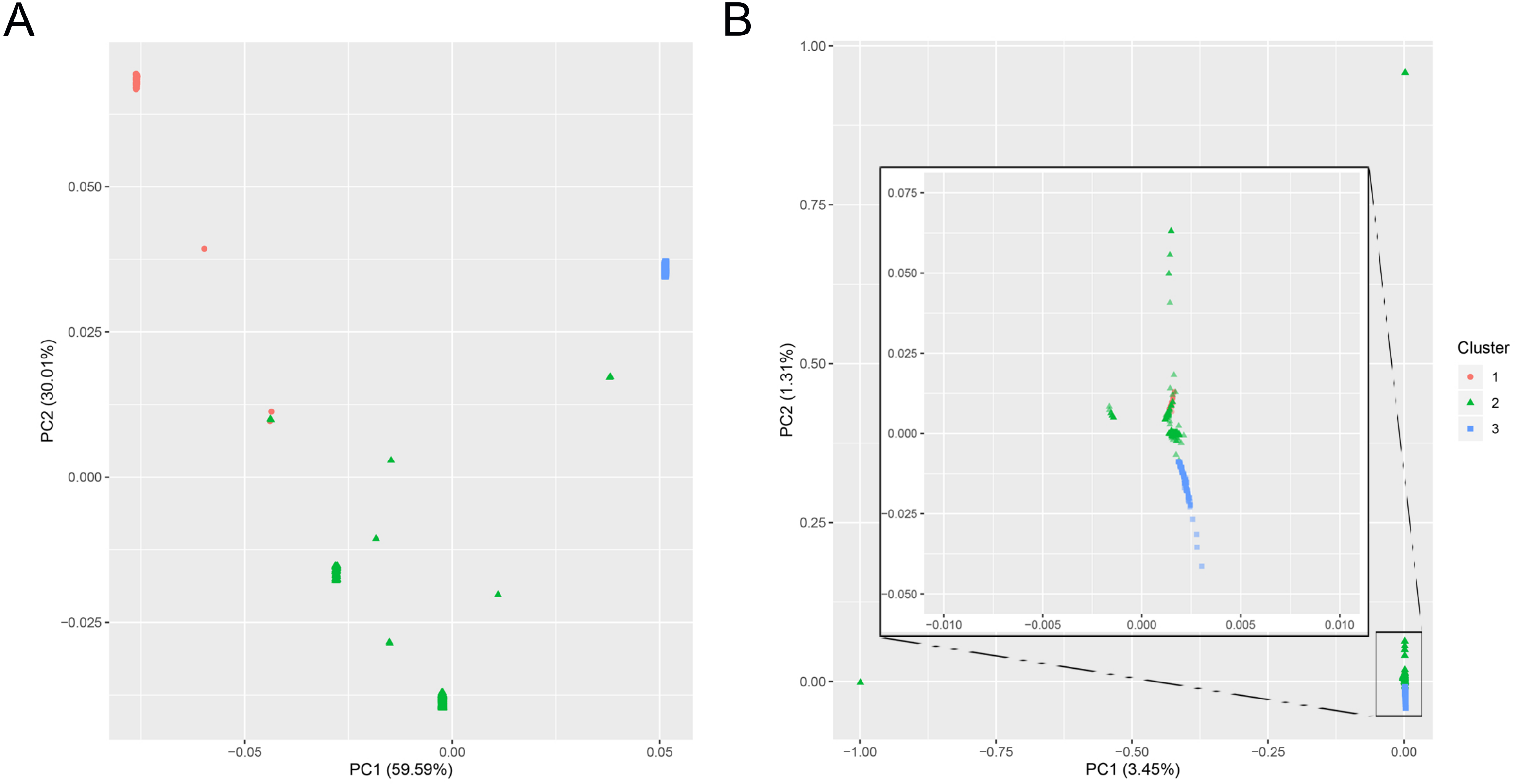
